# NcSnRK1 and NcRbohC1 regulate pollen germination on waterlily stigma after pollination

**DOI:** 10.64898/2026.10.28.762668

**Authors:** Jing Cao, Xin-Rui He, Fan-Rui Min, Chun-Qing Sun

**Affiliations:** Zhenjiang Institute of Agricultural Science in Jiangsu Hilly Areas, Jurong, 212400, China; College of Horticulture, Nanjing Agricultural University, Nanjing, 210095, China

**Author notes:** **Corresponding author: Chun-Qing Sun,**.

**Keywords:** *Nymphaea*, Sexual hybridization, Pollen-stigma recognition, NADPH oxidase, *NcSnRK1*, *NcRbohC1*

## Abstract

Pre-fertilization barrier is the primary cause of intersubgeneric hybridization barrier in waterlily. The research results indicate that reactive oxygen species (ROS) are involved in regulating the germination process of pollen on the stigma of waterlily after pollination, and candidate genes *NcSnRK1* and *NcRbohC1* have been identified; Transient transgenic experiments were used to validate the functions of NcSnRK1 and NcRbohC1 on the waterlily stigma. Through yeast two-hybrid assays, luciferase complementation, and Pull-down experiments, the interaction between NcSnRK1 and NcRbohC1 was confirmed. The above results indicate that NcSnRK1 may phosphorylate certain N-terminal sites of NcRbohC1 after pollination in waterlily, thereby activating ROS burst in the stigmatic papilla cells, disrupting ROS homeostasis on stigma of waterlily, and consequently inhibiting pollen germination on the stigma. The research results can provide theoretical basis for taking effective measures to overcome pre-fertilization barrier and germplasm innovation in waterlily.

## Introduction

Waterlily is a perennial herbaceous flowering plant belonging to the genus *Nymphaea* in the family Nymphaeaceae (Wang et al., 2025). According to the presence or absence of their carpel appendages, waterlily can be divided into subgenus *Nymphaea*, subgenus *Brachyceras*, subgenus *Anecphya*, subgenus *Lotos*, and subgenus *Hydrocallis*. Among them, the subgenus *Nymphaea* is hardy waterlily, while the other four subgenus are tropical waterlily (Sun et al., 2021). At present, cross breeding is still the most common and effective method used in waterlily breeding (Sun et al., 2018; Zhou et al., 2024). However, in the process of cross breeding of waterlily, many breeders have found that intersubgeneric hybridization (hardy waterlilies and tropical waterlilies) has poor seed setting rates, and some hybrid combinations even cannot obtain seeds (Wang et al., 2025; Sun et al., 2019; Sun et al., 2023). This problem greatly affects the efficiency of waterlily breeding and restricts the development and utilization of some excellent color and flower type genes in tropical waterlily. For example, breeders hope to transfer the blue flower gene of tropical waterlily to hardy waterlily in order to obtain blue flower waterlily with cold resistant traits, but most of them have not been successful (Sun et al., 2018). Previous studies have shown that pre-fertilization barrier is the main cause of intersubgeneric hybridization barrier in waterlily (Sun et al., 2019; Hao et al., 2022; Sun et al., 2023). However, the specific reasons for the intersubgeneric hybridization barrier of waterlily have not been reported.

In the early stage of this study, transcriptome and proteomic analysis were conducted on the stigma of waterlily after intersubgeneric hybridization and self-pollination. It was found that ROS was involved in regulating the pollen germination on the stigma of waterlily after pollination, and candidate genes *NcSnRK1* and *NcRbohC1* were identified. This study will focus on the candidate genes *NcSnRK1* and *NcRbohC1*, and conduct in-depth research on the mechanism of intersubgeneric hybridization barrier in waterlily from functional analysis and mechanism of action. It can provide theoretical and technical support for breaking down intersubgeneric hybridization barrier in waterlily and has important guiding significance for cultivating new waterlily varieties.

## Materials and Methods

### Plant materials

Artificial hybridization was carried out using the hardy waterlily (*N*.‘Arc-EN-Ciel’) as the female parent and the tropical waterlily (*N*. ‘Nine Grade Blue’) as the male parent. The parent plants were bagged and isolated one day before flowering. During self-pollination and cross-pollination, the pistils of the female parent plants were emasculated on the day of flowering, and fresh pollen from the male parent plants was injected into the stigma liquid of the female parent plants. After pollination, the flowers were bagged and isolated again. Waterlily plants are preserved in the aquatic flower resource garden of Zhenjiang Academy of Agricultural Sciences in hilly areas of Jiangsu Province.

### Pollen germination on stigma

The germination and growth of pollen on the stigma were observed using the aniline blue staining method. After 6 h pollination, the entire stigma tissue was quickly cut off with a knife and placed in the FAA (70% ethanol : acetic acid: formaldehyde solution = 90 : 5 : 5) fixative solution for more than 12 h of treatment. The fixed stigma tissue was longitudinally cut and soaked in 70% ethanol, 50% ethanol, 30% ethanol, and water in sequence for 10 minutes each, then the stigma tissue was placed in 8 mol/L NaOH solution for softening for 12 h. After softening, it was rinsed with distilled water 3 times and stained with 0.1% aniline blue solution in the dark for 4 h. The pollen tube growth was observed under a upright fluorescence microscope (Ex375-328/DM415/BA351p), and photos were taken using a DSRi2 digital camera (Sun et al., 2015). Each experiment was conducted with three biological replicates.

### Determination of ROS content in stigma

After 0.5 h pollination, the stigma was quickly cut off and immersed in the MES-KCl buffer solution (10 mM MES, 5 mM KCl, 50 mM CaCl_2_, pH 6.15) for 30 minutes. The stigma was then longitudinally cut with a knife, and stained in the dark with 50 mM H_2_DCFDA fluorescent dye for 1.5 h. The sample was then rinsed three times with the MES-KCl buffer solution (Zhang et al., 2021; Wang et al., 2025). After cleaning, the sample was placed on a glass slide and observed and photographed using an upright fluorescence microscope (with the same exposure value of 200) for the analysis of the experimental results.

### Vector construction and transient transgene

The transient gene interference or overexpression of waterlily was achieved by using the improved double virus vector system (IL-60-BS/IR) of tomato yellow leaf curl virus (Ofer et al., 2014; Yuvalet al., 2007). The full-length *NcRbohC1* (*NcSnRK1*) gene was inserted into the PIR-X empty vector to form the PIR-*NcRbohC1* (PIR-*NcSnRK1*) overexpression vector; the conserved region of approximately 300 bp in the *NcRbohC1* (*NcSnRK1*) gene was inserted into the IR-X-RI empty vector to form the IR-*NcRbohC1* (IR-*NcRbohC1*) interference vector. The overexpression vector PIR-*NcRbohC1* (PIR-*NcSnRK1*) or the interference vector IR-*NcRbohC1* (IR-*NcRbohC1*) was mixed with the IL-60-1 plasmid at a ratio of 1:1 and injected into the young leaves of waterlily at a concentration of 800 ng/100 μL, with 200 μL injected each time and the infection performed once a week (Li et al., 2024). Cross-pollination and self-pollination were conducted 14 days after infection. The primers used for gene cloning and vector construction are shown in Table S1, and the gel imaging results of the bacterial liquid detection are shown in Figure S1.

### RNA extraction and real-time quantitative PCR

RNA extraction was performed using FastPure Universal Plant Total RNA Isolation Kit (Vazyme Biotech Co., Ltd; Nanjing); Reverse to cDNA using HiScript III RT SuperMix for qPCR (+gDNA wiper) (Vazyme Biotech Co., Ltd; Nanjing); The ChamQ SYBR qPCR Master Mix was selected as the fluorescent quantitative reagent and measured using the QuantStudioTM real-time fluorescent quantitative PCR system. The relative expression levels of genes were calculated using the ΔCT method, and the Actin11 gene of waterlily was used as an internal reference gene (Luo et al., 2010). The relevant primer designs are shown in Table S1.

### Subcellular localization

For the subcellular localization of proteins in *Nicotiana benthamiana*, the Agrobacterium GV3101 strain carrying pCambia1300-35S:: *NcRbohC1-*GFP (pCambia1300-35S:: *NcSnRK1*-GFP) was cultured at 28°C and 200 rpm/min until OD 600=0.8, and then resuspended until OD 600=1.0. The resuspension was prepared by adding 1 mL 1mol/L MES, 1 ml 1mol/L MgCl_2_ and 100 μL 2 mol/L acetosyringone (AS) to 100 mL distilled water. The resuspension was still in the dark for 2-3 h, and then was immersed into the leaves of *Nicotiana benthamiana* with a sterile syringes. The leaves were cultured overnight in the dark and then cultured under normal light for 48-72 h. The GFP fluorescence signal position was observed using a confocal laser scanning microscope (Leica TCS SP8) to determine the subcellular localization of *NcRbohC1* (*NcSnRK1*).

### Yeast two hybridization

The yeast two-hybrid bait vector pGBKT7-*NcRbohC1*-N (BD-*NcRbohC1*-N) was constructed. Combined with BD-empty and pcl1 plasmids, self-activation verification was conducted respectively in single-missing plate SD/-Trp or SD/-Leu. Using pGADT7 (AD) prey in yeast library and BD-*NcRbohC1*-N, x-α-gal dye was used to screen out the genes that might interact with BD-*NcRbohC1*-N in SD/-Trp-Leu and SD/-Trp-Leu-His-Ade. The gene vector AD-*NcSnRK1*, which may interact with BD-*NcRbohC1*-N, was constructed and verified by yeast one-to-one interaction using x-α-gal dye in SD/-Trp-Leu and SD/-Trp-Leu-His-Ade, respectively.

### Luciferase Complementation Assay

The vectors of pCAMBIA1300-nluc-*NcRbohC1*-N (nluc-*NcRbohC1*-N) and pCAMBIA1300-cluc-*NcSnRK1* (cluc-*NcSnRK1*) were constructed, and the plasmids were transferred into Agrobacterium GV3101 strain with OD 600=1.0. The supernatant was removed by centrifugation, and resuspended to OD 600=0.4-0.6 with the resuspension. After standing at 28℃ in the dark for 3 h, the supernatant was inserted into the leaves of *Nicotiana benthamiana*. After dark culture for 1 d and light culture for 2 d, the luciferase (D-luciferase potassium salt) was used to measure luciferase activity using NightOWL II LB983 in vivo imager (Berthold, Stuttgart, Germany). The suspension solution was prepared according to 2.5.

### Protein induction and pull-down experiments

The *NcRbohC1*-N gene was inserted into the pGEX-4T-1 vector with GST tag, and the *NcSnRK1* gene was inserted into the pCold vector with His tag. The constructed recombinant plasmid was transferred into Escherichia coli BL21 cells to form the *NcRbohC1*-N-GST and *NcSnRK1*-His fusion proteins.

The bacterial liquid was propagated at 37℃ and 200rpm until OD 600=0.6-0.8. 500 μL of bacterial solution was absorbed, centrifuged at 5000 g for 15 min at 4℃, and stored at -80℃ for later use. IPTG was added to the remaining bacterial solution (the final concentration was 0.1 mM) for protein induction. SDS loading buffer was added to 200 uL bacterial solution and then boiled in boiling water at 100℃ for 10 min. SDS-PAGE electrophoresis and protein staining analysis were performed. After successful induction, the protein was centrifuged to obtain the supernatant. The bacterial pellet was resuspended in 1×PBS and separated using an ultrasonic disintegrator (with ice bath throughout, approximately 25 min). After the treatment, the sample was centrifuged at 10000 rpm for 10 minutes at 4℃.

The two proteins that had been ultrasonically broken were mixed and incubated at 4℃ for 2 h in an incubator. 50-100 μL of the product was aspirated and stored at -80℃ for future use. The washed Anti-GST magnetic beads were added to the remaining product and incubated for more than 8 h. The protein magnetic beads that have been incubated were washed with 1×PBS (this process was repeated 3 times). After washing, they were placed in a multi-functional rotating shaker at room temperature for a brief incubation. Then, SDS loading buffer was added, and the mixture was boiled in boiling water for 10 min to denature. Subsequently, polyacrylamide gel electrophoresis and Western blotting were performed.

### Western blot assay

The treated protein samples were separated by 12% polyacrylamide gel electrophoresis for 80 min. The protein glue was imprinted on the PVDF membrane (the PVDF membrane needed to be activated in advance). After transferring the membrane, the membrane was placed into a Petri dish with blocking solution, placed in a shaker at 75 rpm, and blocked at 4℃ for more than 1 h. The membrane was cleaned with 1×PBST for 3 times. The washed membranes were placed in the primary antibody buffer solution, incubated at 75 rpm and 4℃ for 12 h, and then washed 5-6 times. The washed membrane was placed in the secondary antibody buffer, incubated at 75 rpm at 4℃ for 4 h. After washing the membrane 5-6 times again, it was observed using a universal molecular imager.

### Data analysis

All data are presented in the form of mean ± standard deviation. Data analysis was conducted using Microsoft Office Excel. For the analysis of significant differences, one-way analysis of variance was selected, and Tukey’s multiple comparison test or independent sample T-test was used. GraphPad Prism 8 was used for plotting.

## Results

### Pre-fertilization barrier between subgenera

After 6 h of self-pollination of *N*.‘Arc-EN-Ciel’, a large number of pollen tubes germinated and penetrated the stigma disk to reach the ovule for fertilization (Fig. 1A). In the *N*.‘Arc-EN-Ciel’×*N*. ‘Nine Grade Blue’, the pollen grains only adhered to the stigma disk but did not germinate (Fig. 1B).The results indicate that there is a pre-fertilization barrier in intersubgeneric hybridization of waterlily.

**Fig.1.**
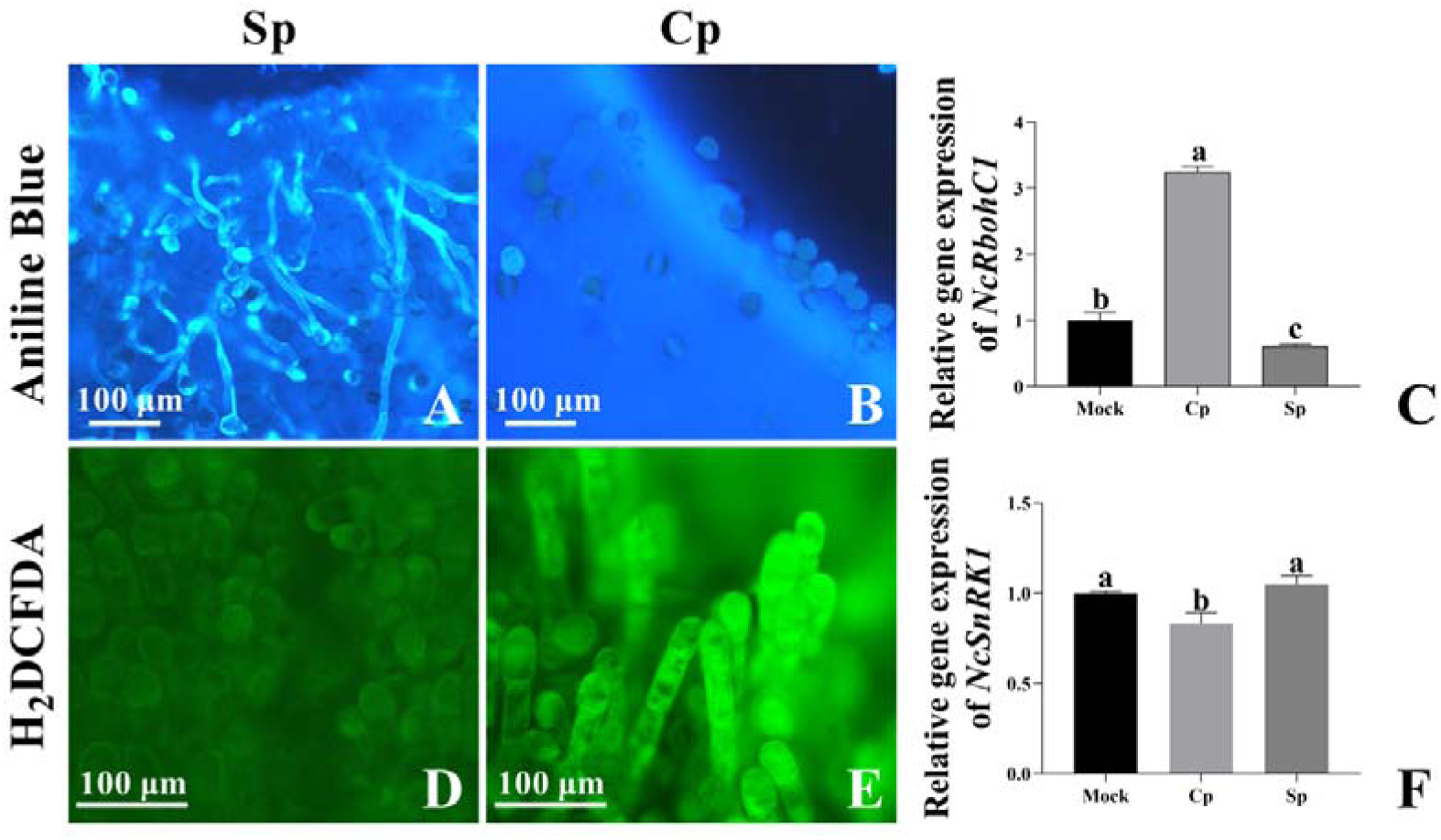
Pollen germination, ROS content and the relative gene expression level of the stigma after cross-pollination and self-pollination of waterlilies. Note: Sp: self-pollination, Cp: cross-pollination, Mock: control.

### ROS is the key regulatory factor

In the preliminary stage of this study, transcriptome and proteomics analyses were conducted on the stigma of waterlily after cross-pollination and self-pollination. It was found that ROS was involved in regulating pollen germination on stigma after pollination in waterlily. The ROS content at 0.5 h in the *N*.‘Arc-EN-Ciel’ × *N*. ‘Nine Grade Blue’ (Fig. 1D) was significantly higher than that in the self-pollination of *N*.‘Arc-EN-Ciel’ (Fig. 1E). This indicates that ROS may play an important role in the regulation of pollen germination on stigma after pollination in waterlily. Half an hour before pollination of *N*.‘Arc-EN-Ciel’ × *N*. ‘Nine Grade Blue’, an antioxidant DMTU (N, N’-Dimethylthiourea) was added to the stigma liquid. The ROS level in the stigma significantly decreased, and a large number of pollen grains germinated on the stigma after cross-pollination (Fig. 2A and D); while half an hour before self-pollination of *N*. ‘Arc-EN-Ciel’, xanthine/xanthine oxidase (X/XO) was added to the stigma liquid. The ROS level in the stigma significantly increased, and it effectively inhibited the germination of pollen on the stigma (Fig. 2B and E). The above experiment demonstrated that ROS is involved in regulating the germination of pollen on the stigma after pollination in waterlily.

**Fig.2.**
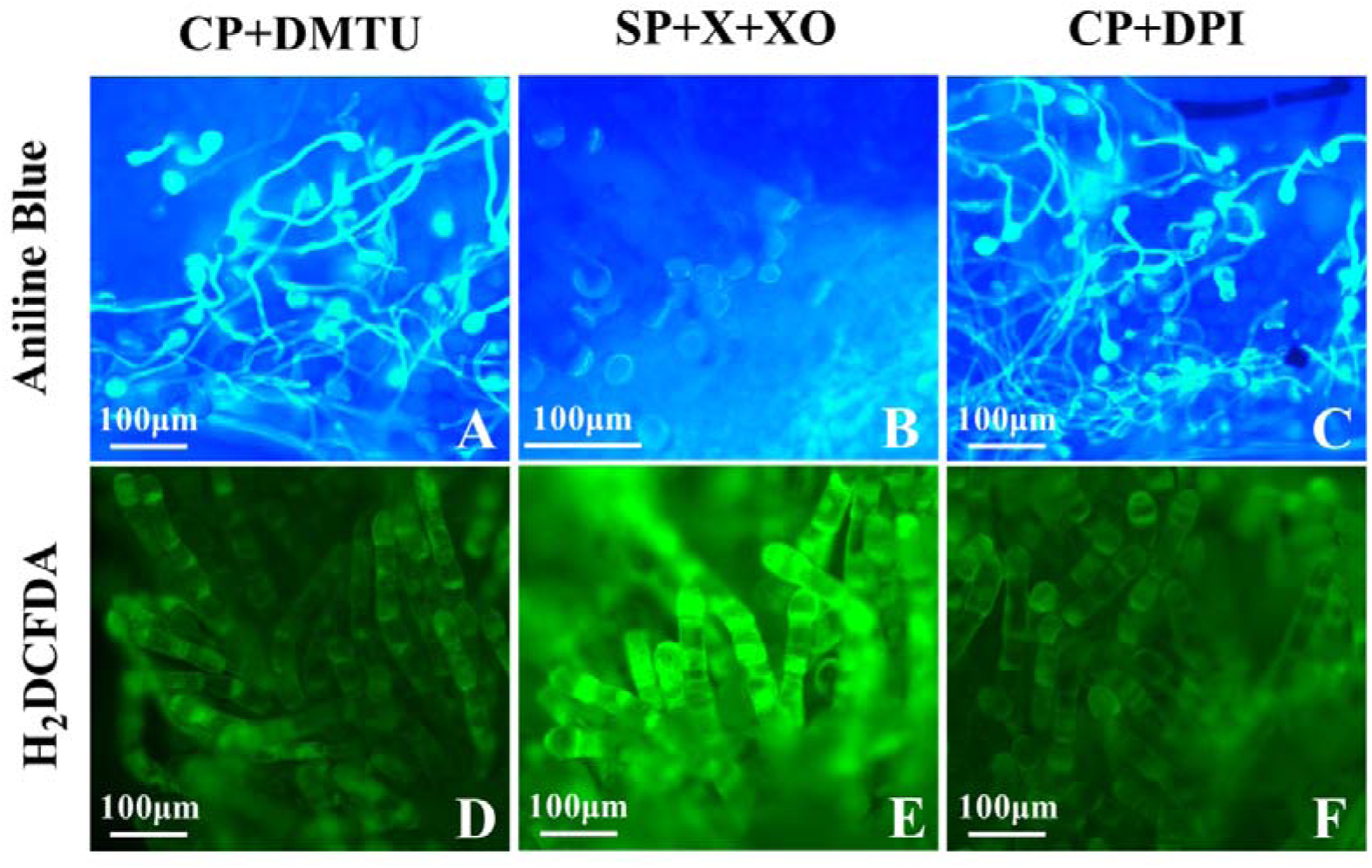
The effects of different active substances on pollen germination and the ROS content in the stigma Note: DMTU: N, N’-Dimethylthiourea; X+XO: Xanthine /Xanthine Oxidase; DPI:Diphenyleneiodonium chloride

### NcRbohC1 associated with ROS generation

Half an hour before pollination, DPI (NADPH oxidase inhibitor) was added to the stigma fluid of *N*.‘Arc-EN-Ciel’ × *N*. ‘Nine Grade Blue’. The ROS level in the stigma decreased significantly, and a large number of pollen germinated on the stigma after cross-pollination (Fig. 2C and F). This indicates that NADPH oxidase is the main source of ROS in the stigma of waterlily. Based on the previous transcriptome data and the verification of stigma gene expression levels after cross-pollination and self-pollination, we found that the expression level of NcRbohC1 showed significant differences (Fig. 1C), suggesting that NcRbohC1 may play an important role in the ROS metabolism process of the stigma of waterlily.

### Functional verification of NcRbohC1

Subcellular localization indicated that NcRbohC1 was located on the cell membrane (Fig. 3). The waterlily plants were treated with interference (IR-*NcRbohC1*) or overexpression (PIR-*NcRbohC1*). After the expression level of NcRbohC1 showed significant changes, cross-pollination and self-pollination were performed (Fig. S2). The qRT-PCR experiment proved that after cross-pollination, the expression level of NcRbohC1 in the stigma of the interference treatment group decreased significantly, while that in the overexpression treatment group increased significantly (Fig. 4A); after self-pollination, the expression levels of the interference and overexpression groups showed opposite trends compared to those after cross-pollination (Fig. 4B). After the transient transgenic treatment for cross-pollination, the pollen of stigma in the interference group germinated and the ROS content decreased significantly (Fig.5A and E), while the over-expressed pollen did not germinate, and the ROS content was high (Fig.5B and F). After transient transgene treatment for self-pollination, the pollen germination quantity in the overexpression group was significantly lower than that in the interference group (Fig. 5C and D), and the ROS content also increased (Fig. 5G-H). Combined with the results of gene function verification (Fig. 4A), it indicates that NcRbohC1 plays an important role in regulating the ROS content in the stigma of waterlily, and NcRbohC1 participates in regulating the germination process of pollen on the stigma after pollination by regulating the ROS content.

**Fig.3.**
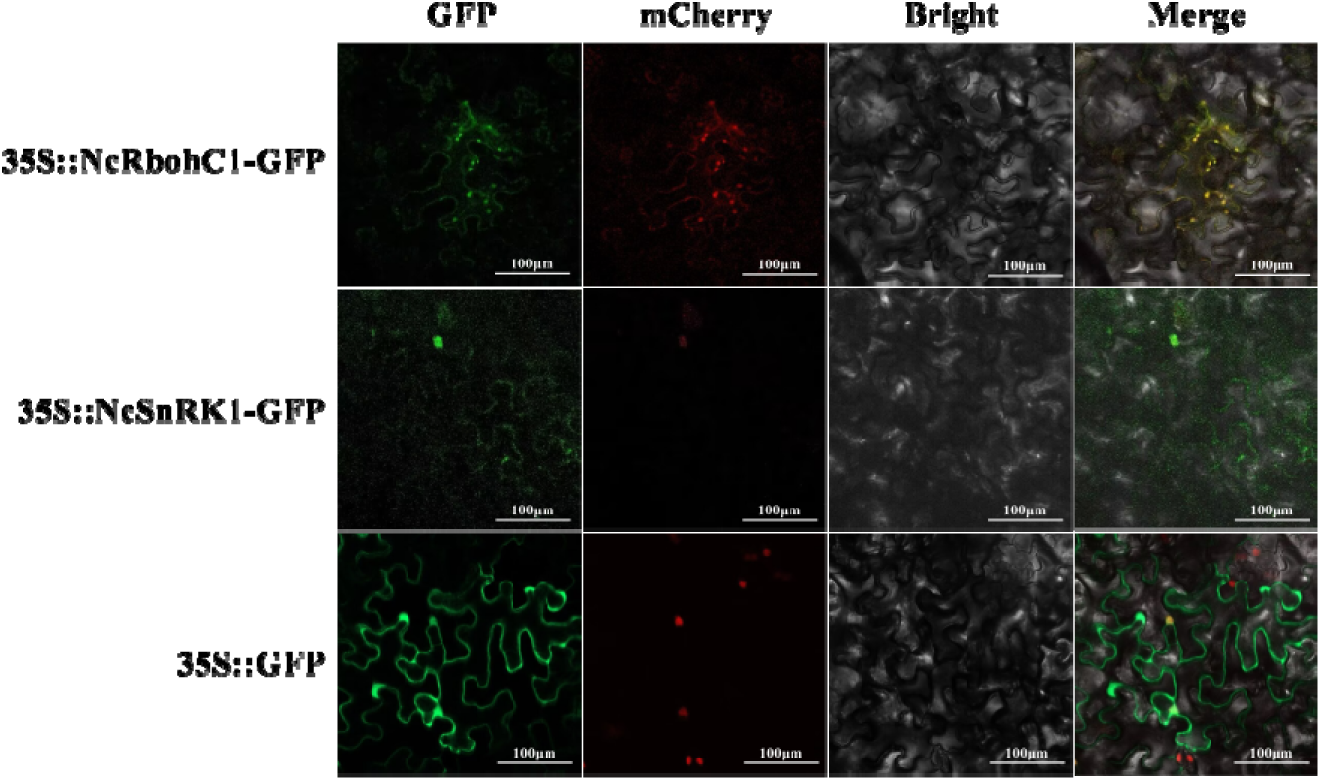
Subcellular localization of NcRbohC1 and NcSnRK1 Note: The subcellular localization of NcRbohC1 used cell membrane marker and NcSnRK1 used cell nucleus marker

**Fig.4.**
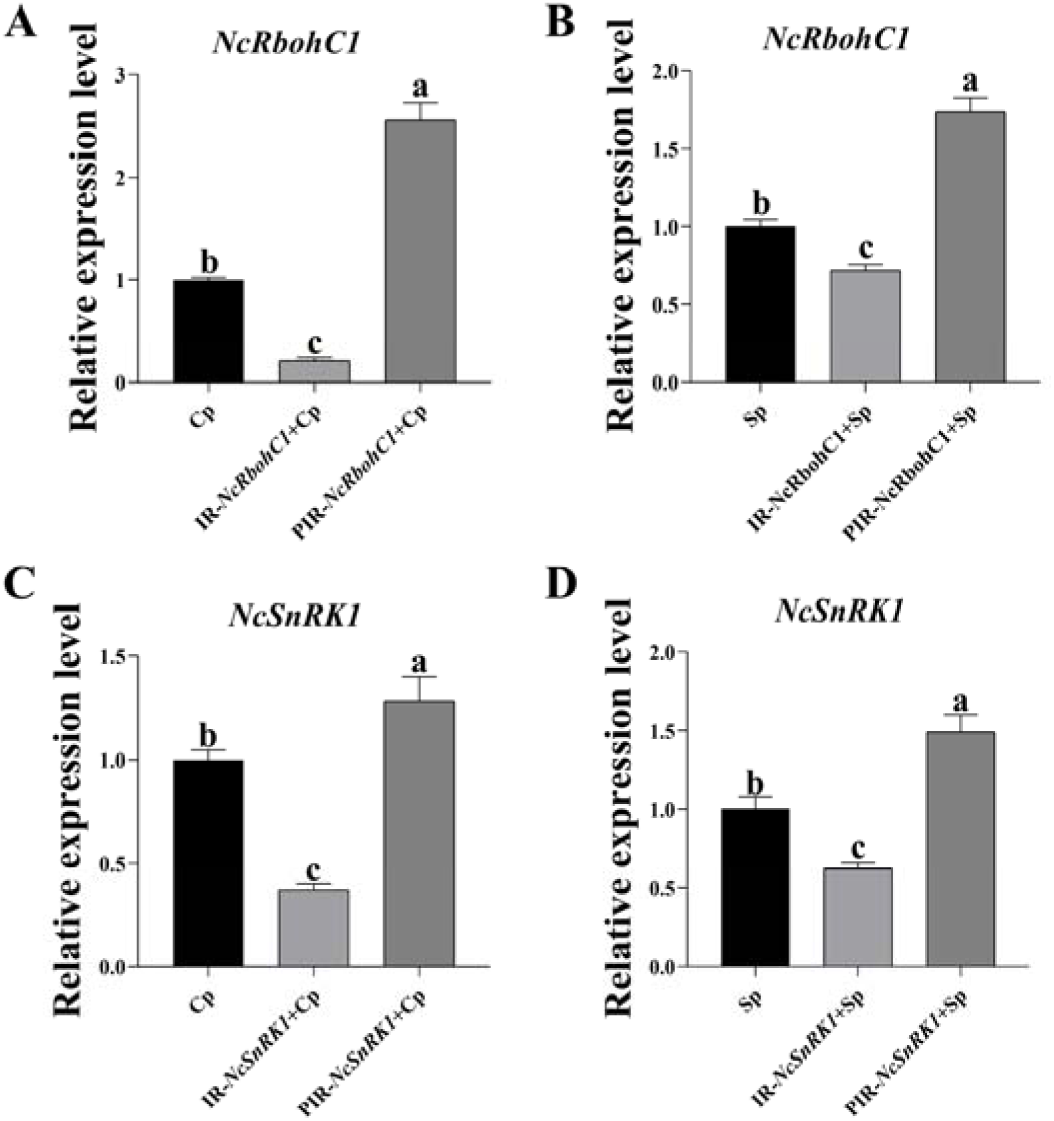
The gene expression level of *NcRbohC1* and *NcSnRK1* after different treatments

**Fig.5.**
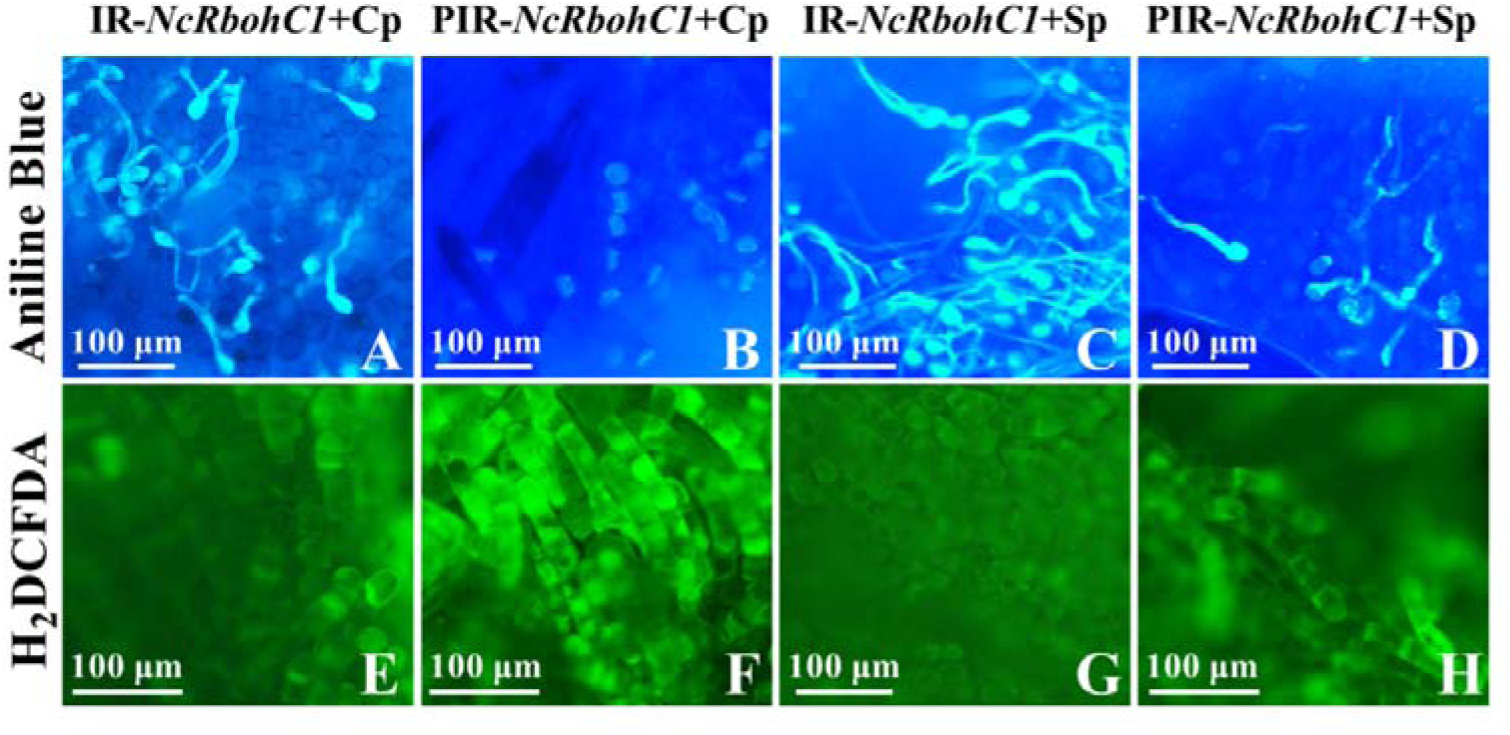
The effects of different treatments on pollen germination and the ROS content in the stigma

### Screening of regulatory proteins of NcRbohC1

To further explore the regulatory mechanism of NcRbohC1 during pollen germination, the pGBKT7-*NcRbohC1*-N (BD-*NcRbohC1*-N) vector was constructed (the N-terminal region of the *NcRbohC1* gene is the functional site), and self-activation verification was performed (Fig. 6A). The existing yeast library was screened. A target gene *NcSnRK1*, which might interact with it and be related to pollen germination, was identified (Fig. S3). The pGADT7-*NcSnRK1* (AD-*NcSnRK1*) vector was constructed, and one-to-one yeast two-hybrid interaction verification experiment was conducted with pGBKT7-*NcRbohC1*-N. The results showed that BD-*NcRbohC1*-N interacted with AD-*NcSnRK1* (Fig. 6B). The pull-down (Fig. 6C) and luciferase complementation (Fig. 6D) experiments also demonstrated that there is an interaction between NcRbohC1-N and NcSnRK1.

**Fig.6.**
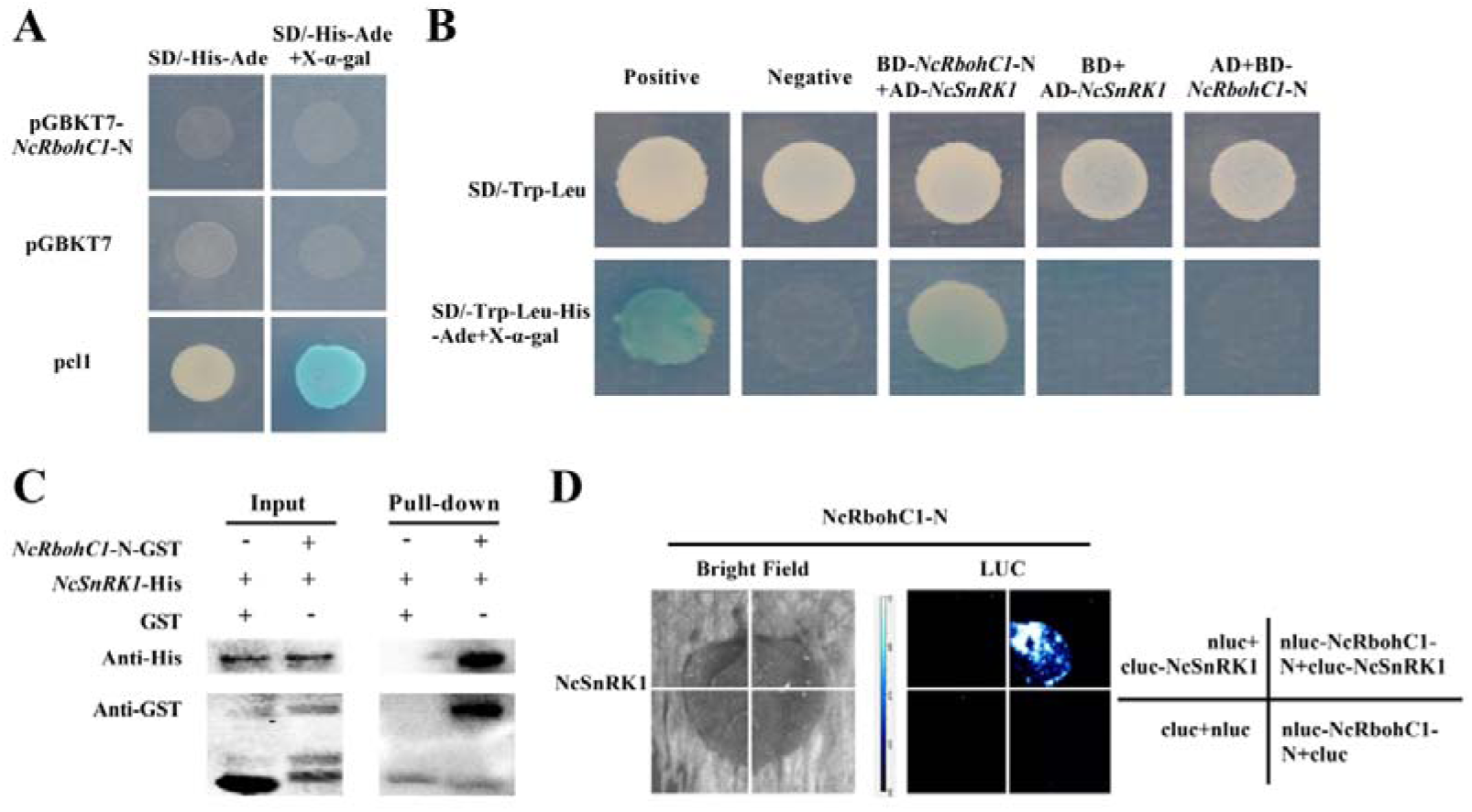
Yeast two-hybrid, pull-down and luciferase complementation assay A: BD-*NcRbohC1*-N yeast self-activation and toxicity detection; B: One-to-one interaction verification; C: pull-down assay; D: Luciferase complementation assay. Note: Positive is the positive control; Negative is the negative control

### Functional verification of *NcSnRK1*

After self-pollination, the expression level of the *NcSnRK1* gene significantly decreased in *N*.‘Arc-EN-Ciel’, but it did not increase significantly after pollination in the *N*.‘Arc-EN-Ciel’ × *N*. ‘Nine Grade Blue’ (Fig. 1F). Subcellular localization indicated that NcSnRK1 was located in the nucleus (Fig. 3). Using transient transgenic technology to treat waterlily plants, the cross-pollination and self-pollination were performed after the expression level of *NcSnRK1* changed (Fig. S4). The results showed that the expression level of NcSnRK1 in the interference group decreased significantly (Fig. 4C), while it increased significantly in the overexpression group (Fig. 4D). Compared with the hybridization after overexpression of *NcSnRK1*, the pollen germination was significantly increased after interference (Fig. 7A and B), and the ROS content was significantly decreased (Fig. 7E and F); while the number of self-pollen germination after overexpression of *NcSnRK1* was significantly lower than that after interference (Fig. 7C and D), and the ROS showed the opposite trend (Fig. 7G and H). This indicates that NcSnRK1 and NcRbohC1 have a synergistic effect on the pollen germination of the stigma.

**Fig.7.**
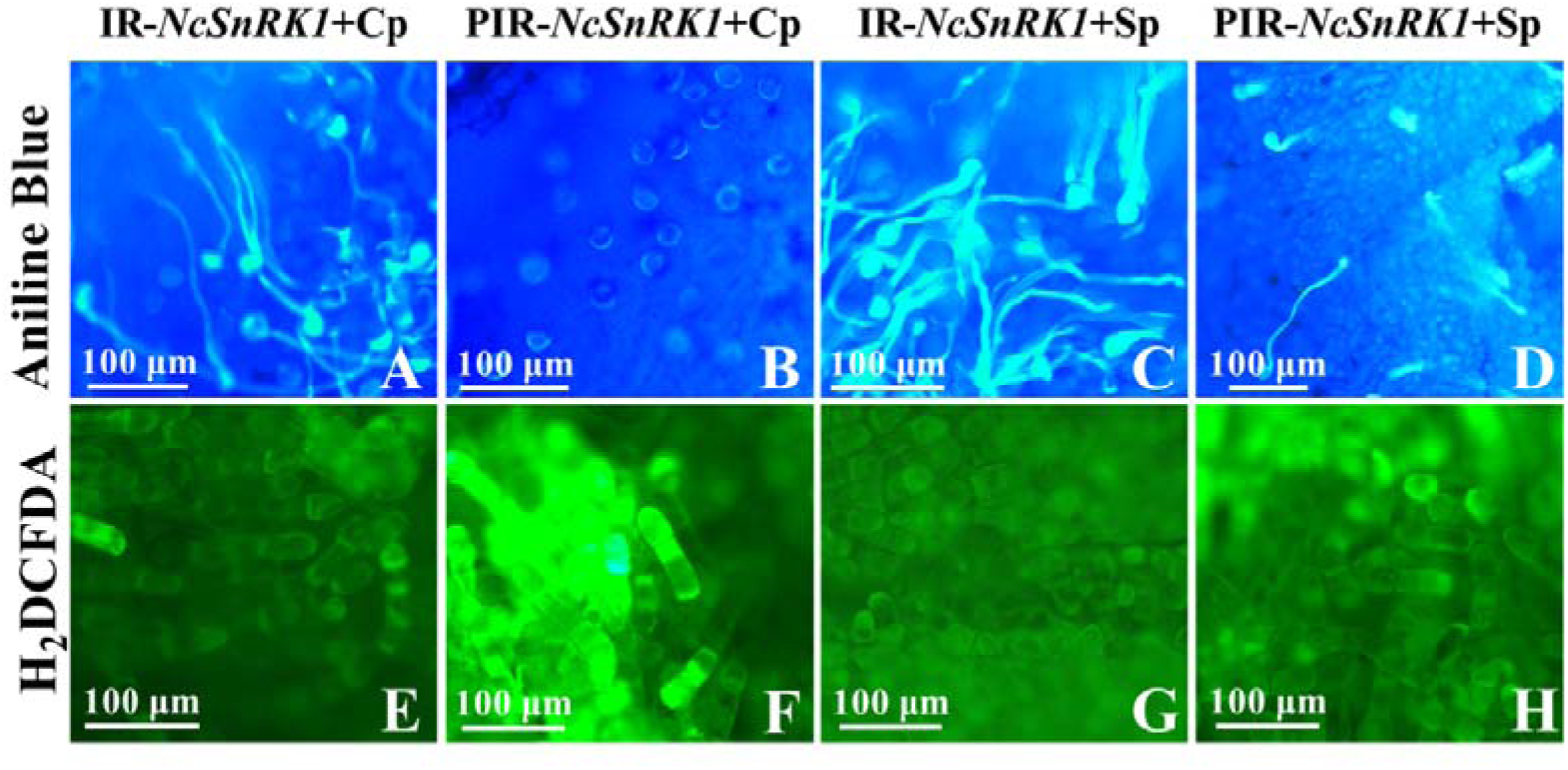
The effects of different treatments on pollen germination and the ROS content in the stigma

### The regulatory relationship between NcRbohC1 and NcSnRK1

Cross-poliination after plant interference/overexpression of *NcSnRK1* had a relatively small effect on the expression level of *NcRbohC1* (Fig.8A), and no significant difference was observed compared with the control group (Cp). Although self-pollination showed a significant difference compared with the control group (Sp) (Fig.8 B), it might be because self-pollination itself can cause changes in ROS content. After overexpression of *NcRbohC1*, both cross-pollination and self-pollination experiments showed a significant increase in *NcSnRK1* expression (Fig. 8 C-D). Conversely, after interfering with *NcRbohC1*, the expression of *NcSnRK1* decreased significantly (Fig. 8 C-D). This indicates that NcSnRK1 may be upstream of NcRbohC1, and may indirectly regulate ROS expression through NcRbohC1.

**Fig.8.**
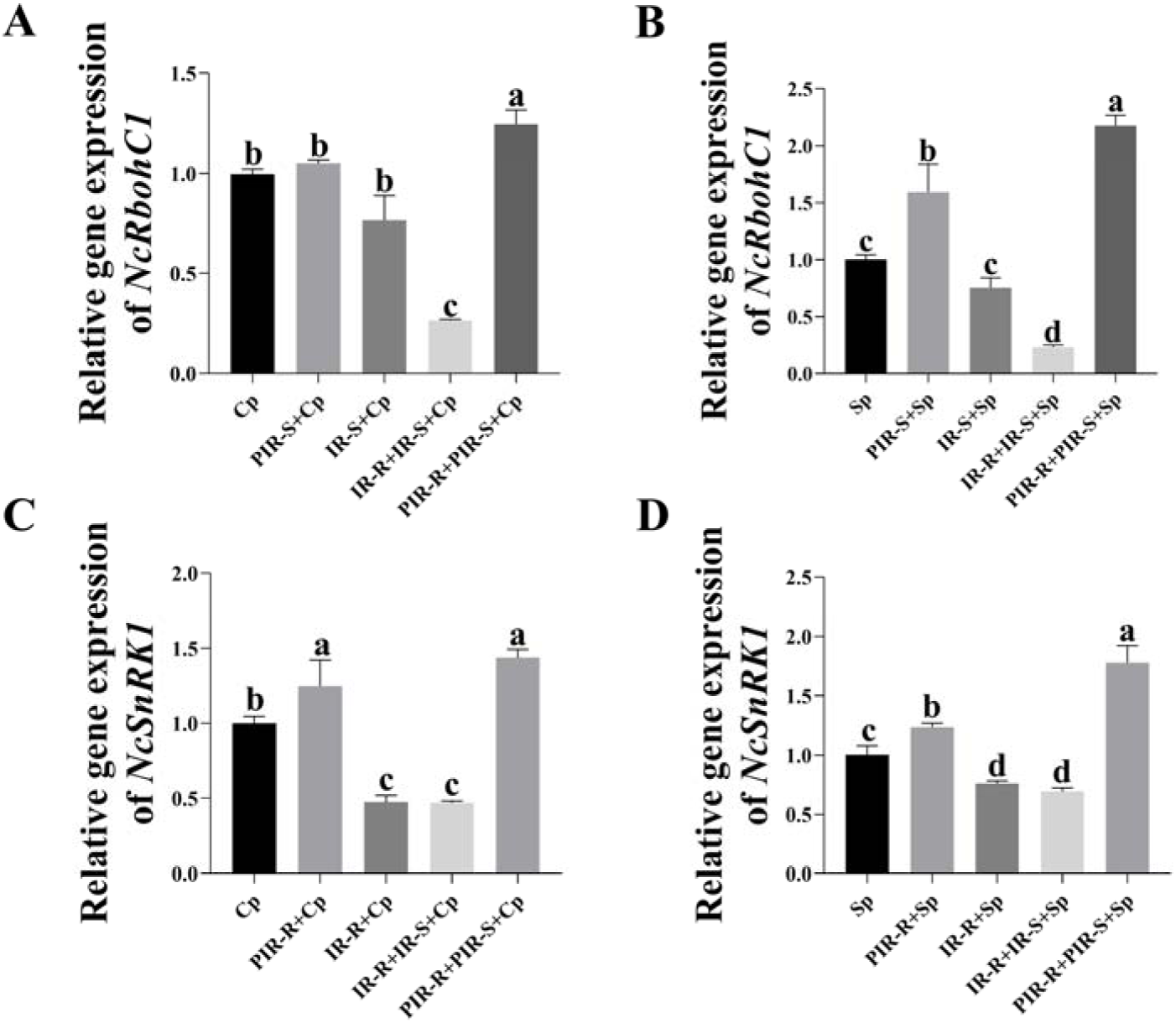
The relative expression level of gene NcRbohC1 and gene NcSnRK1 with different treatments Note: PIR-S:over-expression NcSnRK1;PIR-R:over-expression NcRbohC1;IR-S:interfering NcSnRK1; IR-R:interfering NcRbohC1; IR-R+IR-S:interfering NcRbohC1+interfering NcSnRK1; PIR-R+PIR-S: over-expression NcRbohC1+over-expression NcSnRK1

## Discussion

Most of the existing molecular mechanisms of prefertilization barrier in plant distant hybridization are based on the self-incompatibility (SI) system, which is a typical interspecific hybridization barrier in which the stigma maintains outcrossing by rejecting its own pollen (Huang et al., 2023; Qi et al., 2023). For instance, the SI receptor (S-locus receptor kinase, SRK) on the stigma of Chinese cabbage can recognize self-pollen and activate the ROS pathway on the stigma, increasing the ROS level on the stigma and thereby preventing self-pollen from entering the ovary (Huang et al., 2023; Cao et al., 2025). Similarly, the S-RNase gene expressed in the pistil of pear can recognize the F-box expressed in self-pollen and prevent the pollen tube from extending into the ovary (Qi et al., 2023). Not all plants have the SI system, and the SI systems of plants from different families and genera also vary significantly. Therefore, the molecular mechanism of incompatibility in distant hybridization based on the SI system can only explain some of the hybridization barriers between plant species, and does not have broad applicability.

After the recognition barrier of pollen and stigma, the downstream pathways of stigmatic papillar cells can be further activated, among which the ROS metabolic pathway is more common (Zhang et al., 2021; Huang et al., 2023; Liu et al., 2021.). ROS plays an important regulatory role in self-incompatibility systems of the Papaveraceae (Wilkins et al., 2011; Eaves et al., 2014), *Brassica* (Zhang et al., 2021; Huang et al., 2023), and the genus *Pyrus* (Jiang et al., 2014). Similarly, this study also demonstrates that ROS is involved in regulating the germination of pollen on the stigma after pollination in *Nymphaea*. The NADPH oxidase RBOHs is also known as the respiratory burst oxidase homolog. It is the main source of ROS production in plants. RBOHs have been extensively studied in the process of pollen and stigma recognition (Zhang et al., 2021). This study also demonstrated that NcRbohC1 is a key protein regulating the ROS content in the stigma of waterlily.

ROS production by RBOHs is mainly caused by Calcium-dependent protein kinase (CDPK/CPK) (Kaur and Pati, 2018; Yu et al., 2018), ROP (Rho-related GTPases from plants) protein (Huang et al., 2013), RAC (Ras-related C3 botulinum toxin substrate) protein (Potocký et al., 2012;), etc. Recent studies have found that RBOHs on the stigma are mainly regulated by ROPs to produce ROS. For example, in Chinese cabbage, the FER-RAC/ROPs-RBOHs signaling module can regulate the increase in ROS content on the self-pollination stigma, leading to self-incompatibility (Zhang et al., 2021; Huang et al., 2023). Similarly, in Arabidopsis, the FER/ANJ/LLG1-ROP2-RBOHD signaling module regulates the changes in ROS content (Liu et al., 2021). However, in waterlily, using yeast two-hybrid technology to further screen the interacting proteins of NcRbohC1, it was found that NcRbohC1 does not interact with ROPs, indicating that the regulation mechanism of ROS in the stigma of waterlily is different from that of Chinese cabbage and Arabidopsis. The regulatory mechanism still needs further study. The yeast two-hybrid technique was used to screen the interacting proteins of NcRbohC1. It was found that NcSnRK1 interacted with NcRbohC1. Pull-down and LCA experiments also proved that there was an interaction between NcRbohC1 and NcSnRK1. Moreover, our previous research revealed that NcSnRK1 could regulate the ROS content on the stigma after pollination in waterlily (Zhou et al., 2025). This indicates that NcSnRK1 may regulate the ROS content on the stigma after pollination in waterlily by regulating NcRBOHC1.

The SnRK1 of plants is a homolog of the yeast SNF1 complex and the mammalian AMPK complex (Baena-González and Lunn, 2020). The SnRK1 protein kinase is a heterotrimeric complex composed of the α catalytic subunit and the regulatory subunits β and γ (Baena-González and Lunn, 2020). The function of SnRK1 is mainly determined by the α catalytic subunit, and the function of the α subunit is crucial for understanding the function of the SnRK1 protein kinase (Martinez-Barajas and Coello, 2020). The α subunit contains a kinase domain, and the N-terminal kinase domain and the C-terminal regulatory domain play a key role in the function of SnRK1 (Broeckx et al., 2016; Margalha et al., 2016). SnRK1 has been reported to be involved in regulating the sexual reproduction process of plants. Previous studies have shown that SnRK1 had an impact on the directional growth of pollen, thereby affecting the fertilization process. For instance, in *Arabidopsis thaliana*, the overexpression of SnRK1 inhibits the growth of pollen tubes, thereby affecting fertilization (Baena-Gonzalez et al., 2007). The SnRK1 mutant kinβγ plants of Arabidopsis exhibited pollen abortion, the number of mitochondria and peroxisomes in the pollen was significantly less than that of the wild type, and the level of ROS in the pollen was significantly reduced, which led to the failure of pollen to absorb water and germinate on the stigma (Gao et al., 2016). The SnRK1 mutant kinβγ plants of Arabidopsis exhibited pollen abortion, with significantly fewer mitochondria and peroxisomes in pollen than the wild type, and significantly lower levels of ROS in pollen, resulting inthe failure of pollen to absorb water and germinate on the stigma (Gao et al., 2016). Furthermore, SnRK1 can participate in the regulation of ROS homeostasis. For example, SnRK1α in tomato can phosphorylate serine residues Ser188, Ser189 and Ser308 on RBOH1 protein, and activate RBOH1 to produce apoplasmic ROS (Zheng et al., 2025).

Based on the above analysis, we infer that after cross-pollination among the subgenera of *Nymphaea*, the pollen and stigma encounter recognition barrier, resulting in the extensive expression of NcSnRK1 and NcRbohC1. NcSnRK1 phosphorylates several sites at the N-terminal of NcRBOHC1, thereby activating the ROS burst in the stigmatic papilla cells of the waterlily, leading to the disruption of the ROS homeostasis in the waterlily stigma, and subsequently hindering the germination of pollen on the stigma.

## Author contributions

**Jing Cao:** Investigation, Methodology, Writing – review & editing. **Xin-Rui He:** Data curation, Validation. **Fan-Rui Min:** Data curation, Formal analysis. **Chun-Qing Sun:** Methodology, Supervision, Writing – review & editing.

## Acknowledgments

This study was supported by the National Natural Science Foundation of China (31701948) and the Fund of Independent Innovation of Agricultural Sciences in Jiang Su Province of China [CX(19)3039].

## Abbreviations

ROS: reactive oxygen species
Rboh: Respiratory burst oxidase homolog protein
qRT-PCR: Quantitative Real-time PCR
NADPH: Nicotinamide Adenine Dinucleotide Phosphate Oxidase
DPI: Diphenyliodonium Chloride
DMTU: Dimethylthiourea
X/XO: xanthine/xanthine oxidase
CDPK: Calcium-dependent protein kinase
RAC: Ras-related C3 botulinum toxin substrate
ROP: Rho-related GTPases from plants
SI: Self-incompatibility.

## Figure legends

Fig.S1 The gene cloning of *NcRbohC1, NcRbohC1*-N and *NcSnRK1*

Fig.S2 The gene expression level of *NcRbohC1* after transient transfection

Fig.S3 Yeast two-hybrid screening followed by gel electrophoresis detection

Fig.S4 The gene expression level of *NcSnRK1* after transient transfection Table S1 The primers used in the experiment

